# Ayahuasca Shifts Brain Dynamics Toward Higher Entropy: Persistent Elevation of Ising Temperature Correlates with Acute Subjective Effects

**DOI:** 10.1101/2025.04.25.650509

**Authors:** Rodrigo M. Cabral-Carvalho, Fernanda Palhano-Fontes, Draulio B. Araujo, João R. Sato

## Abstract

Serotonergic psychedelics profoundly alter high-order cognition, emotion, and sensory perception, creating dynamic, unpredictable brain states consistent with the entropic brain hypothesis. Although these acute effects are short-lived, users often experience lasting improvements in emotional and social functioning. Yet, few studies have quantitatively linked subjective experiences to neural dynamics. To bridge this gap, we investigate the acute and subacute effects of ayahuasca using fMRI data and the 2D Ising model, estimating the Ising brain temperature to quantify the shift from ordered to disordered resting state configurations and expose the critical transitions that characterize ayahuasca-induced brain dynamics. To complement our analysis, we investigated cortical functional connectivity changes through graph-theoretical metrics, quantifying how psychedelic ingestion alters functional network architecture in acute and subacute phases. Only the Ising temperature reliably differentiated the subacute phase. Applying a recently developed method that combines cortical connectivity analyses with Graph Neural Networks trained on Ising networks, our results robustly revealed significant Ising temperature increases during the acute (*p <* 0.001) and subacute (*p <* 0.05) phases compared with placebo, reflecting heightened brain entropy and a persistent shift toward a more disordered, paramagnetic state. To evaluate longer-term effects, the subacute temperature change correlated linearly with affective scores on the Hallucinogenic Rating Scale (HRS) captured during the acute session (*p <* 0.01), with HRS scores explaining substantially greater variance in the subacute group (adjusted *R*^2^ = 0.58) than in the placebo group (adjusted *R*^2^ = –0.05). These observations suggest that ayahuasca elevates system entropy and that the magnitude of lasting functional alterations scales with the intensity of the acute subjective experience.

## 1 Introduction

In recent years, scientific interest in the therapeutic potential of serotonergic psychedelics has resurged. These substances primarily influence serotonin receptors, especially the 5-HT2A subtype. Compounds such as psilocybin, lysergic acid diethylamide (LSD), and the traditional Amazonian brew ayahuasca have gained increased attention—not only for their ability to induce profound changes in perception, cognition, and emotion but also for their potential in treating various psychiatric conditions (Maia et al., 2023; Nutt et al., 2020; Palhano-Fontes et al., 2019; Rosenblat et al., 2023). Composed primarily of the leaves of Psychotria viridis and the bark of Banisteriopsis caapi, ayahuasca contains the potent psychedelic molecule N,N-Dimethyltryptamine (DMT) along with reversible monoamine oxidase inhibitors (MAOIs), which together induce a unique and intense altered state of consciousness (Barker, 2018; Fontanilla et al., 2009; McKenna et al., 1984; Riba et al., 2001). Traditionally used by indigenous cultures for spiritual and medicinal purposes, ayahuasca has recently garnered significant scientific interest for its profound effects on consciousness and therapeutic potential (Falchi-Carvalho et al., 2025; Maia et al., 2023; Palhano-Fontes et al., 2019, 2022). Alongside this renewed clinical interest, an expanding body of research seeks to unravel the neurobiological mechanisms underlying the distinctive subjective states and potential therapeutic subacute effects reported following psychedelic experiences.

A central thread in this emerging literature is the exploration of how serotonergic psychedelics impact large-scale brain function. Studies using modern neuroimaging techniques, such as functional magnetic resonance imaging (fMRI), have shown that psychedelics consistently alter patterns of functional connectivity (Carhart-Harris et al., 2012; Palhano-Fontes et al., 2015; Roseman et al., 2014; Viol et al., 2017; Viol Barbosa et al., 2023), particularly within and between canonical networks such as the default mode network (DMN) (Carhart-Harris et al., 2012; Lebedev et al., 2015; Palhano-Fontes et al., 2015; Pasquini et al., 2020). The DMN, closely linked to self-referential processes (Buckner et al., 2008; Raichle et al., 2001), has been shown to become less tightly integrated and more flexible under the influence of psychedelics. This reduction in rigid network patterns can be captured through changes in graph-theoretic measures of connectivity and complexity, suggesting a movement toward more entropic and information-rich brain states (Carhart-Harris, 2018; Viol et al., 2017).

A key framework proposed to account for these functional changes is the entropic brain hypothesis (Carhart-Harris et al., 2014). According to this hypothesis, the subjective intensity and quality of the psychedelic experience correlate with an increase in the entropy, or unpredictability, of neural activity, leading to a temporary shift away from highly constrained, habitual patterns of brain function (Carhart-Harris, 2018; Viol et al., 2019). This expanded repertoire of network states may foster novel insights, emotional processing, and even long-lasting therapeutic “subacute” effects. To investigate these transitions in brain states and entropy gain, studies have used whole-brain models, which integrate empirical neuroimaging data into simulations of large-scale network dynamics, allow systematic manipulation of “control parameters”—for example, global coupling strength, local excitation/inhibition balance, or entropy levels—and then observe how it relates to the resulting patterns of brain activity (Deco et al., 2018; Kringelbach et al., 2020).

Among these models, the two-dimensional (2D) Ising model—commonly used in statistical physics—has proven to be a useful tool for capturing the “temperature” of brain activity, where higher temperature parallels higher entropic states; unlike descriptive entropy measures, Ising temperature provides a single, mechanistic parameter linked to critical phase-transition dynamics that capture the entropy levels of the whole system instead of considering averages (Cabral-Carvalho et al., 2025; Ezaki et al., 2020; Ruffini et al., 2023). By treating the brain as a complex system poised near criticality, we can define an effective “temperature” parameter that reflects the brain’s propensity to fluctuate between ordered and disordered states. Under the influence of psychedelics, increases in this effective temperature may signal heightened neural entropy and complexity, as shown in the acute effects of Lysergic acid diethylamide (LSD) (Ruffini et al., 2023).

Recent studies suggest a potential link between entropy levels and subjective experience, highlighting how the acute phase of psychedelic effects may shape subsequent brain function during the subacute period (Carhart-Harris et al., 2017; Lebedev et al., 2016; Pasquini et al., 2020; Ruffini et al., 2023; Sampedro et al., 2017). No studies until this moment have investigated the changes in temperature for the subacute effects. The present work aims to evaluate Ayahuasca’s acute and subacute impacts on brain dynamics using the 2D Ising Model and how the acute subjective experience relates to lasting functional differences. We also assess functional connectivity with multiple graph metrics, comparing acute and subacute phases to identify overlapping and distinct network alterations. To this end, we analyzed two datasets: (i) task-free fMRI (tf-fMRI) data from nine subjects acquired before and during the acute effects of Ayahuasca and (ii) data from a placebo-controlled trial including tf-fMRI scans acquired before ingestion and 24 hours post-ingestion, along with Hallucinogen Rating Scale (HRS) scores reflecting the acute subjective experience. We hypothesize that Ising temperature will be highest during the acute phase, intermediate during the subacute phase, and lowest under placebo or pre-ingestion conditions. Importantly, we also test whether the Ising temperature measured 24 hours post-ingestion remains elevated and whether this subacute increase correlates with the intensity of the respective acute subjective experience, providing insight into the potential neural basis of psychedelic subacute phenomena. We expect to observe a significant association between Ising temperature and subjective intensity during the acute phase.

## 2 Methods

### 2.1 Data Description

The analyses presented were conducted on two datasets from previous studies, as follows (Figure 1).

**Figure 1:**
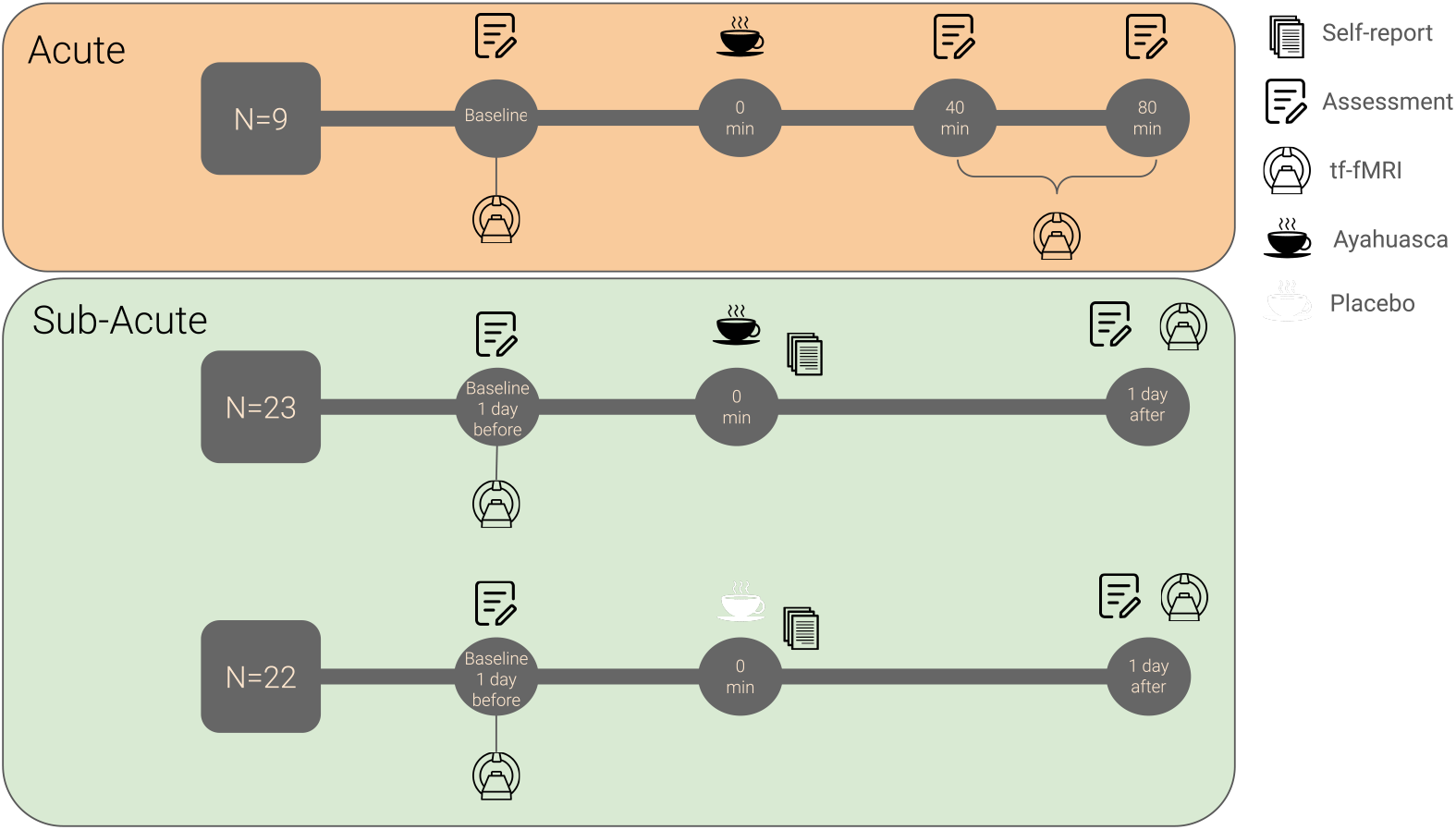
Illustration of the study design, including the acute and subacute ayahuasca experiments, and the fMRI timeline assessments. Black cup: ayahuasca; white cup: placebo.

#### Dataset 1 (acute data)

Nine healthy individuals with at least 5 years of regular (twice a month) ayahuasca use were recruited from the Santo Daime church. Participants underwent two fMRI sessions: one before and another 40 min after ayahuasca intake (2.2 mL/kg of body weight). For more details about the sample, see (de Araujo et al., 2012; Palhano-Fontes et al., 2015). The work was approved by the Ethics and Research Committee of the University of São Paulo at Ribeirão Preto (No. 14672/2006), and written informed consent was obtained from all subjects.

#### Dataset 2 (subacute data)

Forty-three ayahuasca-naïve healthy individuals were randomly assigned to either the ayahuasca group (n = 22) or the placebo group (n = 21). Participants underwent two fMRI sessions: one before and another 24 h after a single dose of 1 mL/kg of ayahuasca or placebo. This dataset is part of a large randomized placebo-controlled trial (Pasquini et al., 2020). All procedures took place at the Onofre Lopes University Hospital, Natal-RN, Brazil. The study was approved by the University Hospital Research Ethics Committee (579.479) and registered at ClinicalTrials.gov (https://clinicaltrials.gov/ct2/show/NCT02914769). All subjects provided written informed consent before participation.

#### 2.1.1 Neuroimaging data acquisition

##### Dataset 1

Images were acquired on a 1.5 T Siemens Magneton Vision scanner. Task-free fMRI data used an EPI sequence (TR = 1700 ms; TE = 66 ms; flip angle = 60°; FOV = 22 cm; matrix = 64×64; slice thickness = 5 mm; 16 slices; 150 volumes). High-resolution T1-weighted images comprised 156 sagittal slices (TR = 9.7 ms; TE = 44 ms; flip angle = 12°; matrix = 256×256; FOV = 25.6 cm; voxel size = 1 mm^3^).

##### Dataset 2

Images were acquired on a 1.5 T GE HDxt scanner. Task-free fMRI data used an EPI sequence (TR = 2000 ms; TE = 35 ms; flip angle = 60°; FOV = 24 cm; matrix = 64×64; slice thickness = 3 mm; gap = 0.3 mm; 35 slices; 213 volumes). T1-weighted images used an FSPGR BRAVO sequence (TR = 12.7 ms; TE = 5.3 ms; flip angle = 60°; FOV = 24 cm; matrix = 320×320; slice thickness = 1 mm; 128 slices; voxel size = 1 mm^3^).

#### 2.1.2 Neuroimaging data preprocessing

Preprocessing and analysis steps were conducted in the same way for both datasets. Initially, the first three fMRI volumes were discarded to allow for T1 saturation. Preprocessing steps were performed using CONN 22 (https://web.conn-toolbox.org/) (Whitfield-Gabrieli & Nieto-Castanon, 2012) standard pipeline, including slice timing correction, head motion correction, and spatial smoothing (6-mm full width at half maximum Gaussian kernel). Functional images were coregistered to the subject’s anatomical image, normalized into the Montreal Neurological Institute template, and resampled to 2*mm*^3^ voxels. The motion artifact was examined using the Artifact Detection Toolbox (ART; http://www.nitrc.org/projects/artifactdetect/). Volumes were considered outliers if the global signal deviation was greater than five SD from the mean signal or the difference in frame displacement between two consecutive volumes exceeded 0.9*mm*. Physiological and other spurious sources of noise were removed using the CompCor method (Behzadi et al., 2007). CompCor is used to reduce non-physiological noise (head movement, cerebrospinal fluid) in BOLD data. Anatomical data define regions of interest composed primarily of white matter and cerebrospinal fluid. Principal components are derived from these noise-related regions of interest and then included as nuisance parameters within general linear models for BOLD fMRI time-series data denoising. Residual head motion parameters (three rotation and three translation parameters), outlier volumes, white matter, and cerebral spinal fluid signals were regressed out. Finally, a temporal band-pass filter of 0.01*Hz* to 0.1*Hz* was applied. We evaluated the amount of head motion in our sample by individually estimating the head mean framewise displacement at both sessions for both datasets. The head motion criteria, specifically head motion *<* 3*mm* translation or *<* 3^*°*^ rotation in any direction, were applied as per previous studies (Ge et al., 2019; Tamm et al., 2006).

### 2.2 Functional Connectivity Measurements

The functional images were segmented into 333 cortical regions of interest (ROIs) based on a parcellation scheme designed to capture functional networks related to cognition, emotion, and sensory processing, with boundaries reflecting consistent connectivity patterns across individuals (Gordon et al., 2016). For each ROI, we extracted the average BOLD signal across its voxels. To quantify functional connectivity, we computed the Pearson correlation between the BOLD signals of different ROIs, forming the adjacency matrix *A* (connectivity matrix). Consequently, the brain network is represented as a graph *G* = (*V, E, A*), where *V* = *{*1, …, *N*} denotes the set of nodes (ROIs), *E* represents the edges, and *A* ∈ ℝ^*N ×N*^ is the weighted adjacency matrix. Each entry *w*_*ij*_ in *A* indicates the connection strength between nodes *i* and *j*, capturing the functional relationship between different brain regions.

To investigate the effects of ayahuasca on brain connectivity, we used several network metrics. Average clustering was used to assess the local grouping of nodes, indicating the extent of community structure and local integration within the network. Average path length provided insights into the efficiency of information transfer across the network, highlighting how quickly and effectively signals can travel between distant regions (McCulloch et al., 2023; Viol et al., 2019). Average betweenness centrality was applied to identify key nodes facilitating communication between different parts of the network, revealing regions relevant for maintaining global connectivity. Additionally, we included segregation, a network granularity measure, given its significance in psychedelics literature (Carhart-Harris et al., 2016). Finally, to further characterize network topology, we reported the standard deviation of degree, reflecting variability in node connectivity, and modularity, assessing the network’s division into distinct functional modules (Daws et al., 2022).

We calculated these metrics across various functional networks defined by Gordon’s Cortical Parcellation that are implicated in cognitive, emotional, and sensory processing: the somatomotor (SMhand and SMmouth), Default Mode Network (DMN), Visual, Fronto-Parietal, Auditory, Cingular Parietal, Retrosplenial Temporal, Cingulo-Opercular, Ventral Attention, Dorsal Attention, and Salience networks (Gordon et al., 2016). These networks were chosen due to their involvement in the altered states of consciousness and self-referential processing observed with ayahuasca use (Palhano-Fontes et al., 2015; Pasquini et al., 2020). This approach enabled us to evaluate both acute and subacute effects of ayahuasca on brain connectivity, focusing on specific local and global alterations that may underlie its cognitive and affective impact.

### 2.3 Ising Temperature

This study employs a Graph Neural Network (GNN) framework to estimate the Ising temperature of functional brain networks derived from resting-state fMRI data developed by (Cabral-Carvalho et al., 2025), using the 2D Ising model as a theoretical foundation for critical dynamics (Fraiman et al., 2009). The 2D Ising model, a statistical physics lattice of interacting spins (1), exhibits phase transitions controlled by temperature (*T*): at low temperatures, the system displays a high degree of order with fewer accessible macrostates (lower entropy), whereas at high temperatures, disorder dominates, increasing the number of accessible macrostates (higher entropy). Near the critical temperature (*T*_*c*_), the system achieves an optimal balance between order and disorder, maximizing the diversity of accessible macrostates and entropy, analogous to hypothesized brain criticality (Chialvo, 2010). Simulated Ising networks across a temperature range were generated using Monte Carlo methods, and their correlation matrices (mimicking functional connectivity) were used to train a GNN to predict the Ising temperature. The trained GNN was then applied to fMRI-derived connectivity graphs (333 ROIs, Pearson correlations) to estimate the brain’s Ising temperature, which can be understood as a measure of global functional connectivity and proximity to criticality. The method enables scalable, computationally efficient analysis of large neuroimaging datasets while bridging theoretical models and empirical brain networks.

To generate the training data for the GNN, we simulated 1.500 Ising networks using Monte Carlo sampling with the Metropolis-Hastings algorithm (Metropolis et al., 1953). The simulations were performed on a 250250 spin-lattice, evolving over 200 time steps after reaching thermal equilibrium. The temperature range spanned *T* = [1.8*J*, 2.5*J*], covering subcritical, critical, and supercritical regimes. The Boltzmann constant and the coupling constant are set to *k* = *J* = 1 so that the temperature has no unit of measurement. The magnetization states were averaged over 13 × 13 blocks, resulting in functional networks with 333 nodes, corresponding to the Gordon parcellation of cortical regions (Gordon et al., 2016). The GNN utilized a graph convolutional architecture, which is well-suited for learning representations from non-Euclidean data structures like brain networks (Kipf & Welling, 2017). More details of the model architecture and training information can be found in Cabral-Carvalho et al., 2025, and the framework can be visualized in Figure 2.

**Figure 2:**
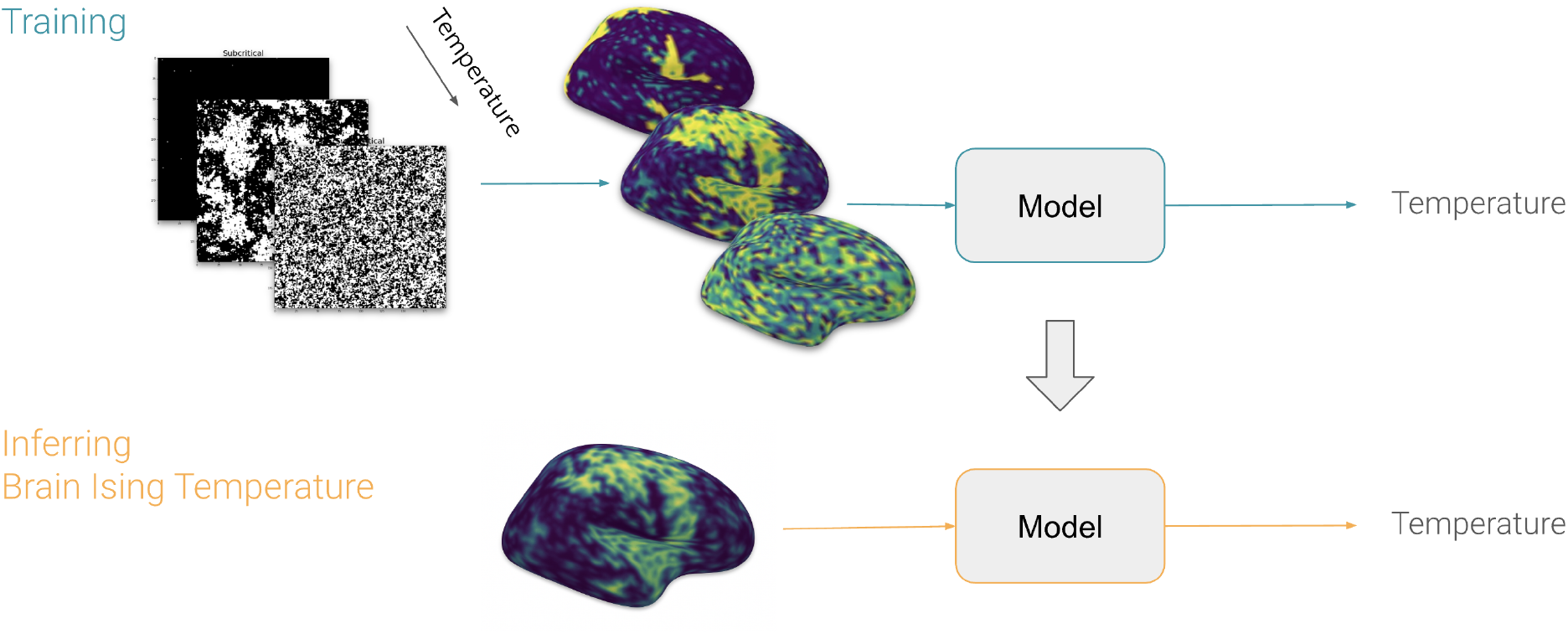
This schematic illustrates the framework for inferring “brain temperature” using 2D Ising models as training data. Multiple 2D Ising configurations generated at different temperatures (left) are fed into a predictive model (top). Once trained, the model is applied to brain data (bottom), yielding an inferred temperature for the brain.

### 2.4 Subjective Effects

For Dataset 2, we analyzed the relationship between brain network measures and acute subjective effects using the Hallucinogenic Rating Scale (HRS) (Mizumoto et al., 2011; Riba et al., 2001; Strassman et al., 1994). The HRS comprises six factors: (i) **Intensity:** overall strength of the experience, (ii)**Somesthesia:** physical sensations (interoception, tactile), (iii) **Affect:** emotional and mood changes, (iv) **Perception:** sensory alterations (visual, auditory), (v) **Cognition:** changes in thought processes, (vi) **Volition:** capacity to control or interact with the experience. Participants completed the HRS approximately four hours post-ingestion of ayahuasca or placebo.

### 2.5 Statistical Testing

The connectivity metrics were analyzed to detect alterations in brain network organization, including changes in local connectivity within brain regions and global connectivity across the brain. Changes in functional connectivity and Ising temperature were assessed: for datasets 1 and 2, scores were calculated by subtracting baseline values from post-treatment values for each participant under the ayahuasca and placebo conditions. These scores in functional connectivity and Ising temperature were then compared using paired t-tests to evaluate whether the magnitude of change differed between conditions. Cohen’s d was calculated to quantify the effect size of these differences.

Additionally, we used linear regression to test whether head-motion variables explain any changes in Ising temperature. All p-values were corrected for multiple comparisons using false discovery rate (FDR) adjustment with the Benjamini–Hochberg method across all network measurements. For the subacute group, we conducted linear regression to assess whether the HRS subscales’ mean scores and standard deviations predict changes in Ising temperature. In this analysis, HRS subscale scores served as independent variables, whereas Ising temperature was the dependent variable (see Figure 3. We focused on statistically significant coefficients indicating a predictive relationship between the HRS metrics and Ising temperature. This approach allowed us to assess how different dimensions of subjective experience relate to changes in brain dynamics. All the p-values were corrected for the testing within networks. The significance level was set to *α* = 0.05, two-tailed. All analyses were performed using SciPy 1.14.

**Figure 3:**
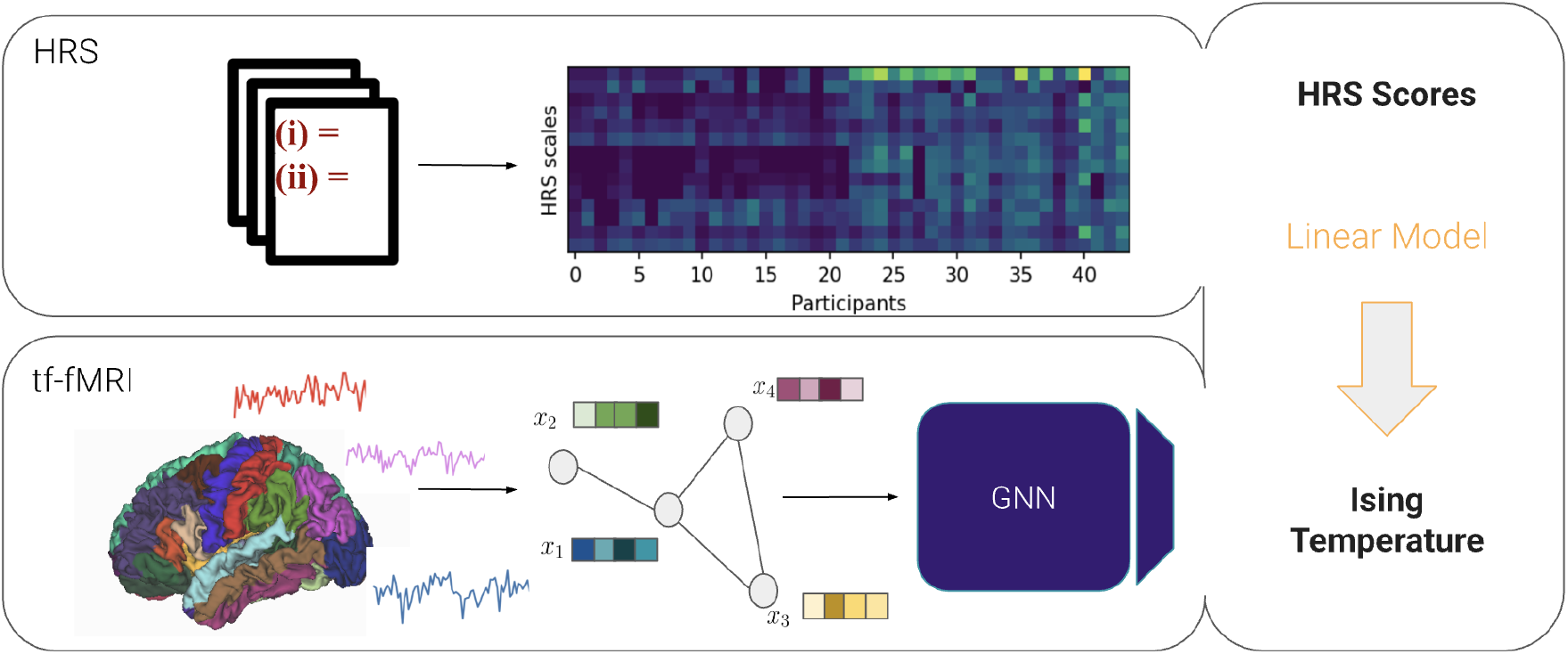
Workflow Diagram to connect Ising temperature from functional connectivity to subjective measures. tf-fMRI data are analyzed through a GNN model to estimate Ising temperature using functional connectivity (Cabral-Carvalho et al., 2025).

## 3 Results

For the final analysis, participants who deviated from resting-state conditions—either lacking a restingstate scan or exhibiting head motion beyond the threshold established in the Methods—were excluded. Specifically, one subject was excluded from the acute dataset (*N* = 8), and four subjects were excluded from each group in the subacute dataset (*N*_placebo_ = 18, *N*_ayahuasca_ = 19).

### 3.1 Functional Connectivity

#### 3.1.1 Local connectivity

During the acute phase, several significant local network changes were identified (Table 1; Figure 4). Metrics exhibiting only decreases included segregation, average betweenness centrality, modularity, and degree standard deviation. Segregation significantly decreased in the Default Mode Network, Auditory, Cingulo-Opercular, and Dorsal Attention networks. Average betweenness centrality decreased in the Somatomotor Hand, Auditory, Cingulo-Opercular, and Ventral Attention networks. Modularity decreased in the Somatomotor Hand, Auditory, and Ventral Attention networks, while the Auditory network also showed increased degree variability.

**Table 1:** Comparison of local brain network metrics (segregation, average clustering, average betweenness centrality, average path length, modularity and degree standard deviation) during the acute phase following ayahuasca administration, showing significant alterations in the Default Mode, Somatomotor Hand, Auditory, Cingulo Opercular, Ventral Attentional, and Dorsal Attentional networks between the acute ayahuasca and baseline. Only metrics with *p <* 0.05 are shown.

| Network | Metric | $t$ | $p$ | $p_{\text{corr}}$ | Cohen's $d$ |
| --- | --- | --- | --- | --- | --- |
| Default | segregation | 4.7 | $> 0.01$ | 0.01 | -1.65 |
| SMhand | average clustering | -4.1 | $> 0.01$ | $> 0.05$ | 1.44 |
| SMhand | average betweenness centrality | 2.8 | $> 0.05$ | 0.05 | -0.97 |
| SMhand | average path length | -3.6 | $> 0.01$ | $> 0.05$ | 1.26 |
| SMhand | modularity | 2.6 | $> 0.05$ | 0.05 | -0.93 |
| Auditory | segregation | 2.8 | $> 0.05$ | $> 0.05$ | -0.98 |
| Auditory | average clustering | -5.6 | $> 0.001$ | $> 0.01$ | 1.96 |
| Auditory | average betweenness centrality | 4.4 | $> 0.01$ | $> 0.01$ | -1.53 |
| Auditory | average path length | -5.3 | $= 0.001$ | $> 0.01$ | 1.87 |
| Auditory | degree SD | 3.2 | $> 0.05$ | $> 0.05$ | -1.19 |
| Auditory | modularity | 3.2 | $= 0.01$ | $> 0.05$ | -1.14 |
| CinguloOperc | segregation | 5.7 | $> 0.001$ | $> 0.01$ | -2.03 |
| CinguloOperc | average betweenness centrality | 4.1 | $> 0.01$ | $> 0.01$ | -1.44 |
| CinguloOperc | average path length | -3.5 | $= 0.01$ | $> 0.05$ | 1.22 |
| VentralAttn | average clustering | -4.1 | $> 0.01$ | $> 0.05$ | 1.45 |
| VentralAttn | average betweenness centrality | 3.1 | $> 0.05$ | $> 0.05$ | -1.08 |
| VentralAttn | average path length | -3.4 | $= 0.01$ | $> 0.05$ | 1.21 |
| VentralAttn | modularity | 3.2 | $> 0.05$ | $> 0.05$ | -1.14 |
| DorsalAttn | segregation | 4.2 | $> 0.01$ | $> 0.05$ | -1.50 |

**Figure 4:**
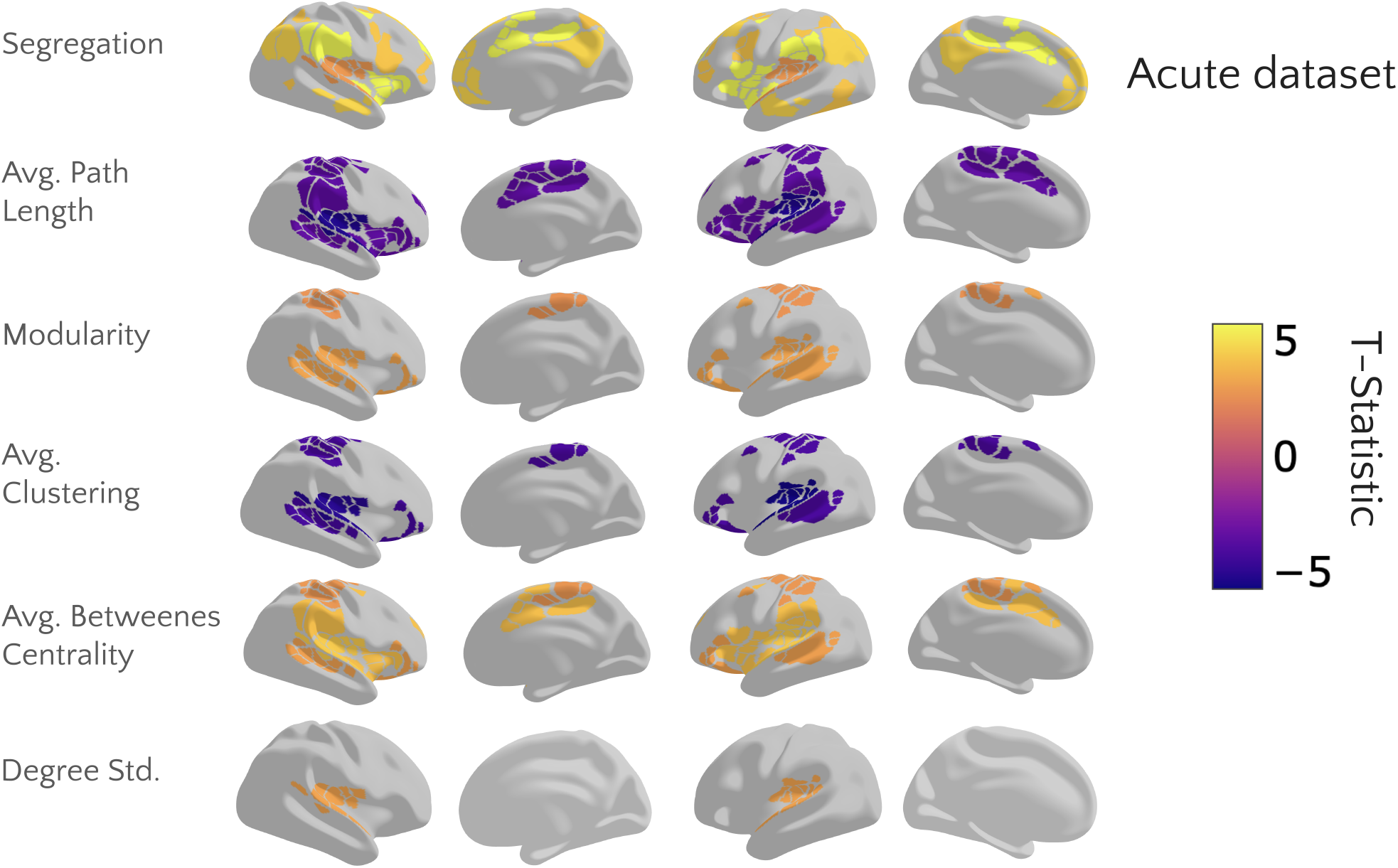
Visualization of brain network metrics in the acute dataset, indicating significant changes across various graph-theoretical metrics applied to fMRI data. The metrics illustrated include segregation, average clustering, average betweenness centrality, average path length, modularity, and degree standard deviation. Each panel maps these changes across the brain, with the color scale representing the t-statistic of these changes, only statistically significant, where warmer colors (yellow to red) denote lower t-statistic, suggesting higher statistical significance, and cooler colors (purple) indicate higher t-statistics, denoting less significant changes. These visualizations capture the statistically significant areas where ayahuasca influences brain connectivity and complexity.

Conversely, metrics exhibiting only increases included average clustering coefficient and average path length. Significant increases in average clustering were observed in the Auditory, Ventral Attention, and Somatomotor Hand networks. Average path length increased in the Somatomotor Hand, Auditory, Cingulo-Opercular, and Ventral Attention networks. These results underscore the complex alterations in network dynamics during the acute phase of ayahuasca ingestion.

Changes in connectivity during the subacute phase can be visualized in Table 2 and Figure 5. There was a decrease in segregation within the Ventral Attentional Network in the ayahuasca group. The Dorsal Attention network was also modified by an increase in average clustering and average degree, although most differences of the subacute group did not survive the multiple comparisons correction.

**Table 2:** Summary of changes in local brain network metrics (segregation, average clustering, average betweenness centrality, average path length, modularity, and degree standard deviation) after ayahuasca administration, highlighting significant differences between the subacute ayahuasca and placebo groups. Only metrics with *p <* 0.05 are shown.

| Network | Metric | $t$ | $p$ | $p_{\text{corr}}$ | Cohen's $d$ |
| --- | --- | --- | --- | --- | --- |
| SMmouth | average clustering | -2.6 | = 0.01 | > 0.05 | 0.81 |
| SMmouth | average betweenness centrality | 2.9 | > 0.01 | > 0.05 | -0.90 |
| SMmouth | average path length | -2.8 | > 0.01 | > 0.05 | 0.86 |
| SMmouth | modularity | 2.6 | > 0.05 | > 0.05 | -0.78 |
| Visual | average clustering | -2.6 | = 0.01 | > 0.05 | 0.80 |
| Visual | average path length | -2.7 | = 0.01 | > 0.05 | 0.82 |

**Figure 5:**
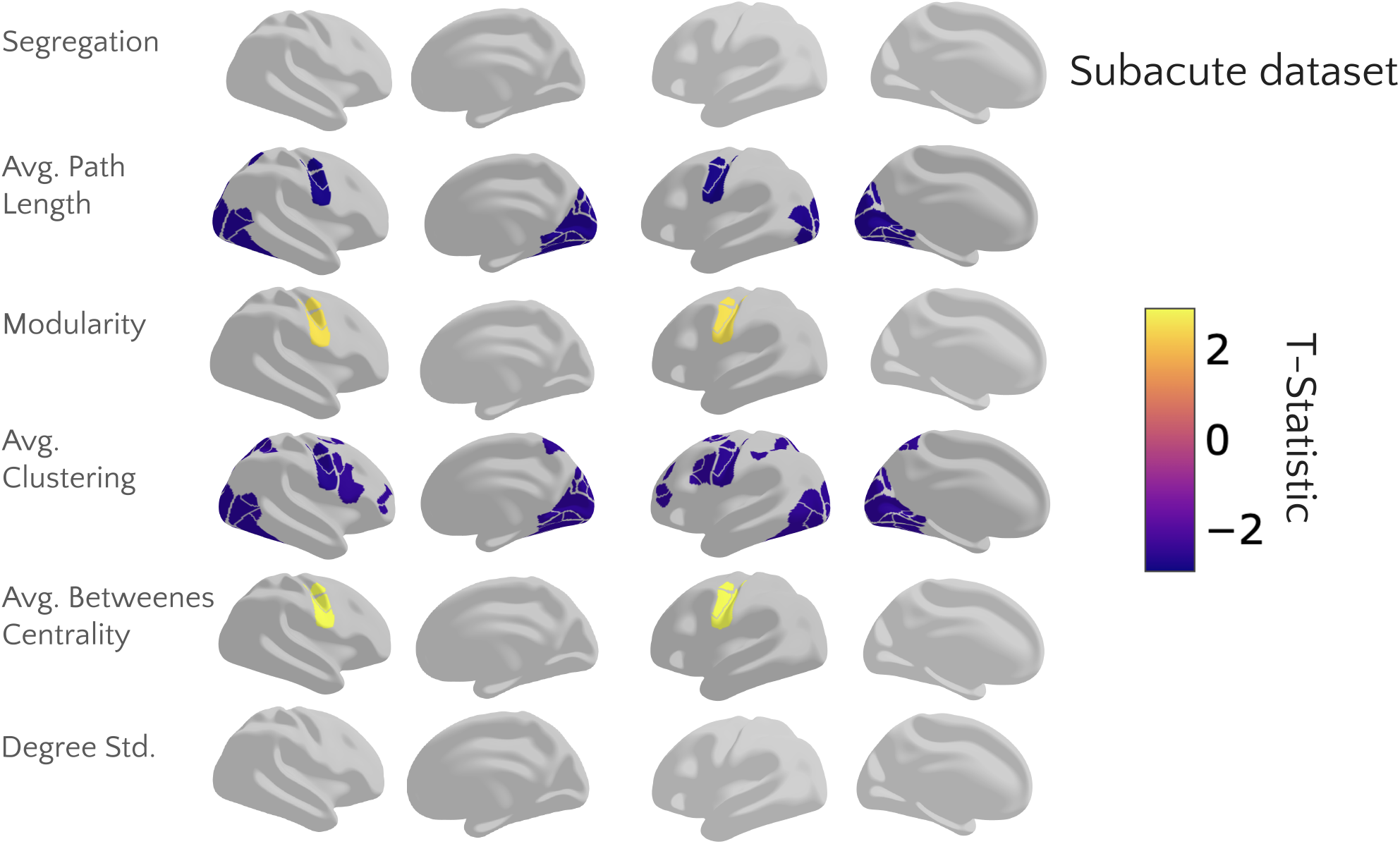
Visualization of brain network metrics in the subacute dataset, indicating significant changes across various graph-theoretical metrics applied to fMRI data. The metrics illustrated include segregation, average clustering, average betweenness centrality, average path length, modularity, and degree standard deviation. Each panel maps these changes across the brain, with the color scale representing the t-statistic of these changes, only statistically significant, where warmer colors (yellow to red) denote lower t-statistic, suggesting higher statistical significance, and cooler colors (purple) indicate higher t-statistic, denoting less significant changes. All p-values were corrected for multiple comparisons using the false discovery rate (FDR) correction with the Benjamini-Hochberg method to control for false positives. These visualizations capture the statistically significant areas where ayahuasca influences brain connectivity and complexity.

#### 3.1.2 Global connectivity

In the acute dataset, although three metrics showed uncorrected *p <* 0.05, only average clustering coefficient and average degree remained significant after FDR correction (see Supplementary Table S1). Specifically, both average clustering and average degree differed between acute ayahuasca and baseline (*p* = 0.02 for both), indicating increased local connectivity and overall network integration under ayahuasca.

In the subacute dataset, although some metrics—such as average path length and clustering—initially showed *p <* 0.05, none survived correction for multiple comparisons. Consequently, these findings were not reported in the supplementary materials. Similarly, in the placebo group, no network metrics exhibited statistically significant changes after correction (all corrected *p >* 0.1), and results were likewise omitted.

### 3.2 Ising Temperature

Our methodology interprets Ising temperature as a measure of connectivity, enabling us to examine whether it correlates with any of the graph metrics used in this study. Our analysis indicates no significant correlations between Ising temperature and global graph metrics during acute or subacute phases. The Pearson correlation coefficients for these metrics were all below 0.25, with associated p-values exceeding the conventional threshold for statistical significance (*p >* 0.1), as detailed in the Supplementary material (Table S2). During the acute effects, all participants exhibited an increase in Ising temperature, whereas in the subacute effects, only a few participants showed an increase. For individual data, refer to the supplementary material (Figure S1 and Figure S2, respectively).

For the subacute dataset, before drug administration, no significant differences were observed between the groups (ayahuasca vs. placebo) (*p* = 0.2), suggesting comparable baseline brain states. Post-administration, the ayahuasca group exhibited a notable increase in |*T* − *T*_*c*_| relative to the placebo group (p = 0.01). To assess the differences between the MRI scans for the acute ayahuasca and subacute and placebo groups, we computed the changes in Ising temperature (*δT* = *T*_*after*_ − *T*_*baseline*_). The analysis revealed a significant difference between the subacute and placebo groups (*p* = 0.02) and between the acute and baseline (*p <* 0.001). However, no statistical difference was found in temperature between the acute and subacute datasets, as illustrated in Figure 6.

**Figure 6:**
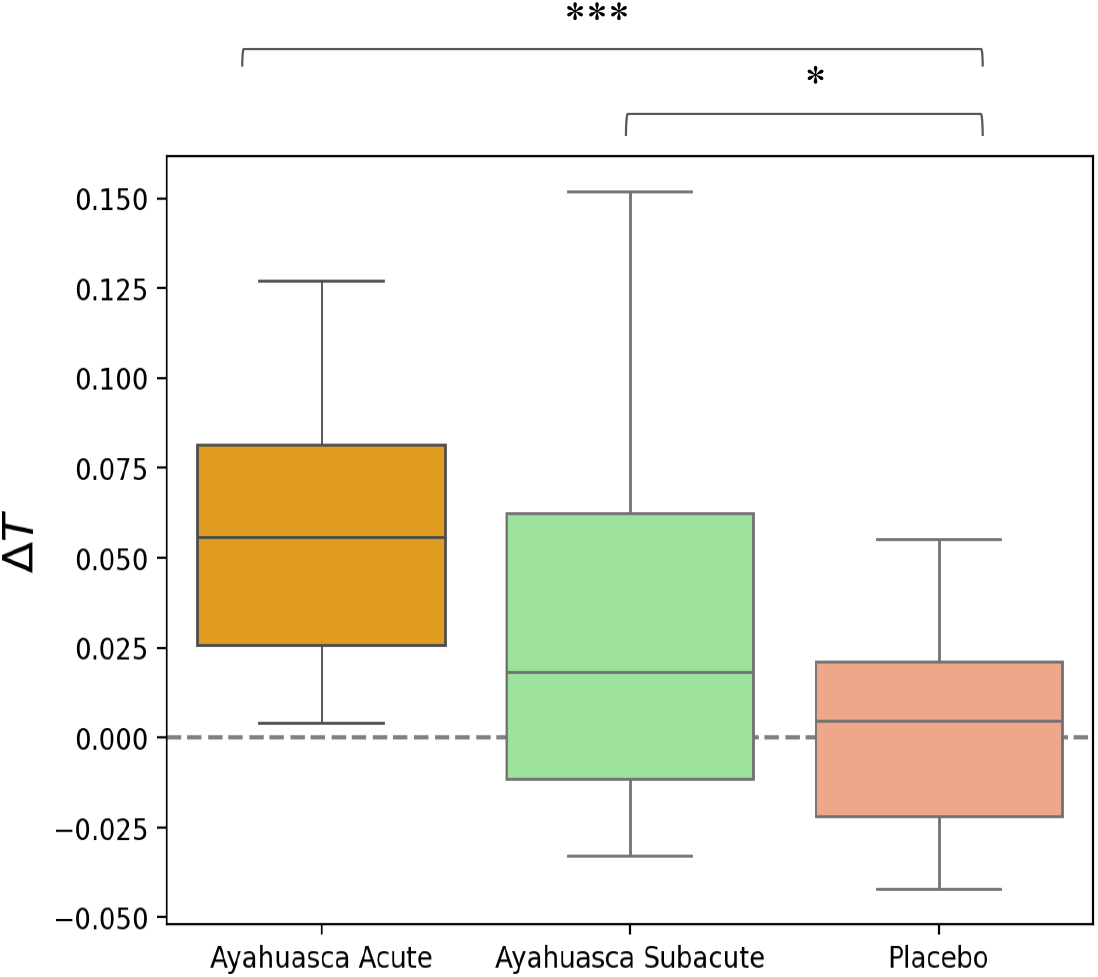
Box plot comparing changes in Ising temperature (*δT*) between the placebo, ayahuasca subacute, and ayahuasca acute groups. The plot shows that the median *δT* is higher in both the subacute and acute phases following ayahuasca administration compared to the placebo group, indicating increased brain entropy and complexity. The variability in *δT* suggests distinct alterations in cortical functional connectivity induced by ayahuasca across different time points. Asterisks denote significance (\**p <* 0.05, * * *p <* 0.01, * \*\**p <* 0.001).

#### 3.2.1 Confounding analysis (Motion)

To assess the influence of potential confounds, the impact of head motion on Ising Temperature estimation was examined (see Table S3), given its significant effect on fMRI data (Friston et al., 1996). A linear regression analysis determined which variables could predict the Ising Temperature. The independent variables included Translation in X, Y, and Z (in mm), and Rotation in X, Y, and Z (in degrees), with the Estimated Ising Temperature as the dependent variable for each group. While the acute dataset moved their heads more than the subacute, the results indicate that none of the head motion parameters significantly impacted the Ising Temperature estimates in the acute or subacute datasets, with all p-values exceeding the threshold for statistical significance. These findings highlight that head motion does not significantly influence Ising Temperature estimates and appears to be a consistent predictor across different groups.

### 3.3 Linking function to subjective experience

For the subacute group, a linear regression analysis, where the independent variables were the HRS subscale scores and the dependent variable was the Ising temperature, was conducted to assess whether the average scores and standard deviations of the HRS subscales can predict the increase in Ising temperature among subjects (See Figure 7, which shows t-values from the linear regression predicting the Ising Temperature based on the HRS subscales for the placebo and ayahuasca groups, for details see Table S4). By examining the coefficients and t-statistics, we observe that the average affect score (*p <* 0.01) and the standard deviation of the cognition score (*p* = 0.04) significantly contribute to explaining variations in temperature increases. Notably, the adjusted *R*^2^ = 0.58 for the ayahuasca group and adjusted *R*^2^ = −0.05 for the placebo group, indicating that, for placebo participants, the HRS subscales did not meaningfully predict Ising temperature changes.

**Figure 7:**
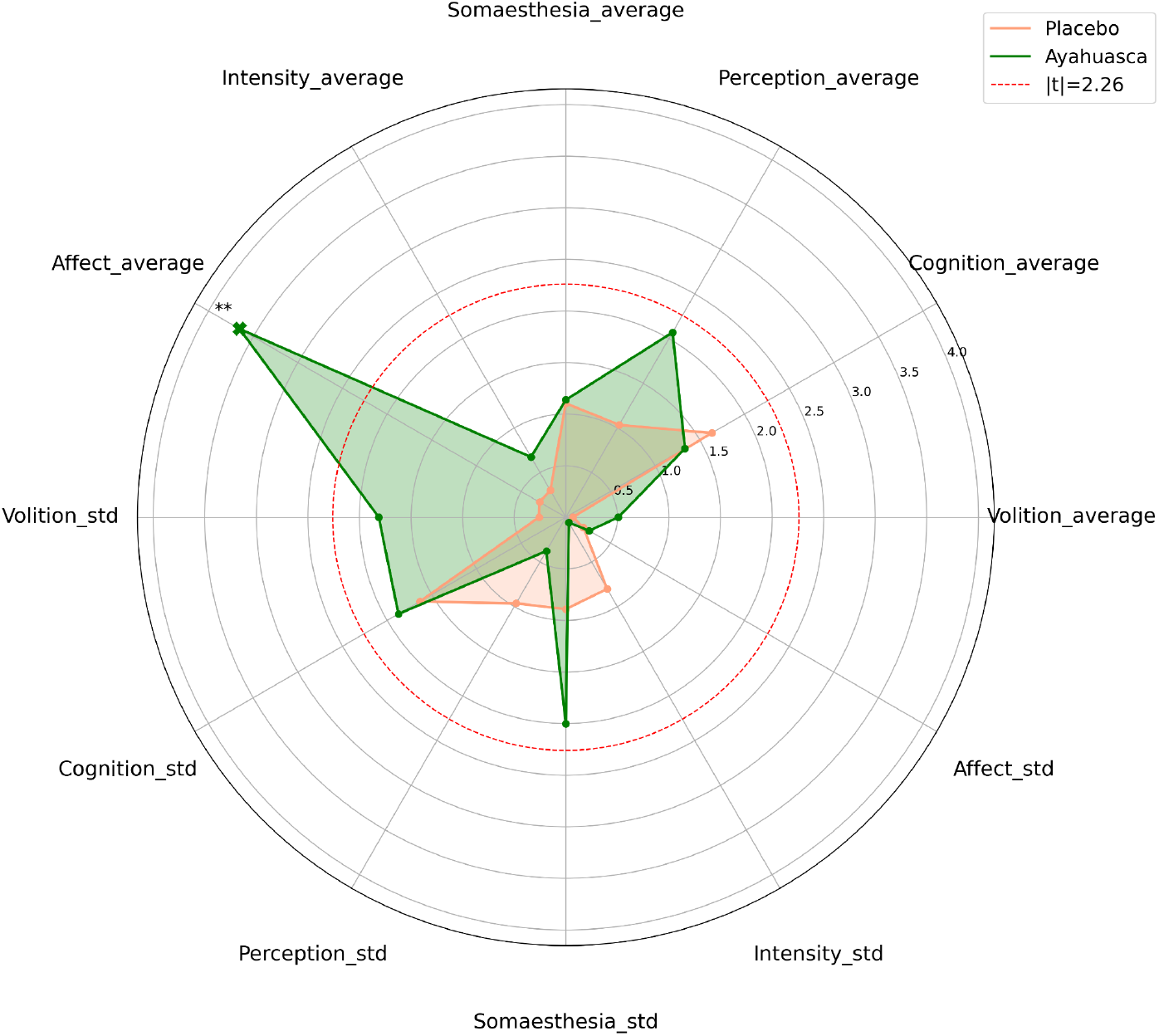
Radar chart displaying the t-values from a linear regression predicting the Ising Temperature based on the Hallucinogenic Rating Scale (HRS) subscales for the placebo (orange) and ayahuasca (green) groups. Each spoke represents a different HRS dimension—intensity, somaesthesia, affect, perception, cognition, and volition—in terms of mean (average) and variability (standard deviation). The dotted red circle marks *t* = 2.26—a use threshold for statistical significance at *p* = 0.05 given nine residual degrees of freedom. The visualization shows all the t-statistics in absolute value, and the marker demonstrates the sign of the significant variables (positive “+” and negative “-”). The plot illustrates how ayahuasca produces significantly different contributions (in multiple subscales) to the model predicting Ising Temperature compared with placebo. (* *p <* 0.05, ** *p <* 0.01).

## 4 Discussion

In this investigation, we assessed the acute and subacute neural effects of ayahuasca by measuring brain Ising temperature and connectivity metrics through rs-fMRI data. Consistent with prior research, we found reduced network segregation and modularity during the acute phase (Palhano-Fontes et al., 2015; Pasquini et al., 2020), a phenomenon widely observed across psychedelic states (Carhart-Harris et al., 2012; Gattuso et al., 2023). This breakdown of canonical community structure appears to support a more globally integrated brain state, potentially underlying core psychedelic effects such as ego-dissolution, sensory blending, and increased cognitive flexibility (Carhart-Harris et al., 2016; Tagliazucchi et al., 2016). We also observed increased betweenness centrality within the Ventral Attention and Cingulo-Opercular networks, identifying them as possible integration hubs, as well as increases in average path length and average clustering, markers of more exploratory yet local cohesive dynamics (McCulloch et al., 2023; Viol et al., 2017, 2019). Despite the decrease in modularity the increase in local average clustering, shows tight local triangles, perhaps around newly emergent hubs. As segregation and modularity are key features of network complexity, our findings suggest that ayahuasca acutely reorganizes functional connectivity, possibly facilitating novel integration pathways through specialized hubs, similar to effects reported for psilocybin, LSD, and DMT (Daws et al., 2022; Roseman et al., 2014; Timmermann et al., 2023). Notably, we observed no overlap in networks showing significant changes in clustering and path length across the acute and subacute phases, indicating distinct modes of network reconfiguration over time.

Apart from classical connectivity measures, we also evaluated the Ising temperature through functional connectivity as a measure of brain dynamics. The results revealed significant increased Ising temperature during the acute phase across all subjects, indicating heightened entropy and a paramagnetic shift, along with a sustained elevation in temperature during the ayahuasca subacute in some individuals. Despite the fundamentally different approach to infer the Ising temperature (Ezaki et al., 2020), the results were consistent with previous findings on the acute effects of LSD, in which the brain shifts toward a more disordered, paramagnetic state, as also reported by Ruffini et al., 2023. However, this study is the first to extend such analysis to both the subacute phase of psychedelics and to ayahuasca specifically. This reflects enhanced neurodynamic flexibility induced by ayahuasca. The rise in Ising temperature corresponds to increased availability of macrostates, greater disorder, and reduced predictability in neural dynamics. Notably, the subacute group exhibited statistically significant increased Ising temperature compared to placebo. Ising temperature captures variations in predictability and the number of accessible brain states through a physical model, monotonically relating to the system entropy. Thus, our approach closely aligns with the entropic brain hypothesis (Carhart-Harris et al., 2014; Ruffini et al., 2023), which posits that psychedelic substances enhance brain entropy, expanding the spectrum of cognitive states and amplifying subjective experiences characteristic of psychedelic states.

Most interestingly, the subacute Ising temperature change was considerably better explained by HRS subscale scores compared to placebo differences, highlighting that subjective experiences during the acute phase may significantly influence subacute effects on brain dynamics. Explicitly correlating with scores on the affect subscale of the Hallucinogen Rating Scale (HRS). This correlation in affect scores aligns with previous research linking affective shifts to altered brain connectivity in subacute effects of ayahuasca (Pasquini et al., 2020). Interestingly, the relationship between neural changes and subjective experiences was uniquely analyzed in the subacute phase due to the absence of HRS in the acute group, revealing notable individual variability. In this later period, only certain participants maintained elevated Ising temperatures, indicating a selective persistence of heightened entropy and reduced neural predictability. The connection between the HRS and the persistent rise in system entropy emphasize that subjective experiences during the acute phase may significantly influence subacute effects on brain dynamics. This might suggest that acute-phase experiences could be crucial in driving sustained neurobiological and psychological transformations. These findings offer evidence of ayahuasca’s lasting and profound impact on brain function, providing compelling insights into its effects on consciousness, emotional states, cognitive flexibility, and therapeutic potential.

The experimental design presents further limitations. The acute dataset had a small sample size and lacked randomization or placebo control. There were differences between the acute (see details in Palhano-Fontes et al., 2015) and subacute (see details in Pasquini et al., 2020) datasets, including dose, scanner type, and participant background. Finally, while Ising temperature offers a useful abstraction, its direct biological meaning remains exploratory. The use of Ising temperature as a proxy for neural activity relies on a highly simplified representation of brain dynamics. Although the 2D Ising Model captures fundamental emergent phenomena, it significantly simplifies the complexity and nuanced neurophysiological processes occurring during psychedelic experiences (Fraiman et al., 2009). Furthermore, this model’s limitations extend to specific assumptions inherent in our current machine-learning-based framework: it employs a constant coupling matrix for all spins, restricts interactions exclusively to immediate neighbors, omits the external magnetic field, and assumes the chosen model accurately reflects empirical brain data (Cabral-Carvalho et al., 2025); however, the results agree with other methods to infer Ising temperature (Ezaki et al., 2020; Ruffini et al., 2023). Additionally, our analysis based predominantly on average connectivity metrics may overlook critical aspects of brain networks. Metrics focused only on averages can mask important variations, distributions, spread, and entropy within network connections. A deeper exploration of these distributional properties might uncover subtle but significant differences in brain network behavior induced by psychedelics, enhancing the interpretability and precision of results.

Recognizing and addressing these limitations is essential for developing more accurate and comprehensive models of complex neural interactions.

## Data and Code Availability

All analysis code and supplementary materials are available at https://github.com/Rodrigo-Motta/Ayahuasca.

## Author Contributions

Conceptualization: R.C.; Data curation: R.C, F.P.; Methodology: R.C.; Investigation: R.C.; Resources: R.C., F.P., D.A.; Formal analysis: R.C.; Visualization: R.C.; Writing—original draft: R.C.; Writing—review & editing: R.C., F.P., J.S., D.A.; Project administration: D.A., J.R.; Software: R.C.; Supervision: D.A., J.S.; Funding acquisition: R.C., J.S., D.A.

## Acknowledgements

This work was supported by São Paulo Research Foundation (FAPESP) Grants 2021/05332-8, 2023/02616-0, 2023/02538-0; Brazilian National Council for Scientific and Technological Development (CNPq) grants 466760/2014 & 479466/2013; and CAPES Foundation grants 1677/2012 & 1577/2013.

