## Supplementary Material for "Ayahuasca Shifts Brain Dynamics Toward Higher Entropy: Persistent Elevation of Ising Temperature Correlates with Acute Subjective Effects"

| Metric | t_statistic | p_value | p_corrected |
|---|---|---|---|
| average_clustering | -3.6 | > 0.01 | > 0.05 |
| average_betweenness_centrality | 1.8 | < 0.05 | > 0.05 |
| average_path_length | -2.3 | = 0.05 | > 0.05 |
| std_degree | -3.9 | > 0.01 | > 0.05 |
| modularity | 3.0 | > 0.05 | > 0.05 |

**Table S1:** Overview of global brain network metric changes in the acute dataset (segregation, average clustering, average betweenness centrality, average path length, modularity and degree standard deviation) observed during the acute phase after ayahuasca administration, indicating significant alterations in network-wide connectivity patterns between the ayahuasca and placebo groups.

**Relation between Ising temperature and connectivity**

We fitted a multiple linear regression model using Ordinary Least Squares (OLS), with the overall change as the dependent variable and the individual metric changes as predictors. In order to facilitate the interpretation of each predictor's contribution, we standardized both the independent variables and the dependent variable using z-score normalization. This standardization allowed the regression coefficients to be directly compared in terms of their relative effect sizes.

The standardized model produced coefficients that indicate the number of standard deviations the dependent variable is expected to change per one standard deviation change in the predictor. To estimate the relative contribution of each predictor to the explained variance, we squared each standardized coefficient and computed its proportion relative to the sum of all squared coefficients. Expressing these proportions as percentages provided a quantitative measure of the importance of each metric in explaining the overall change.

This approach not only yielded a robust model for predicting change but also offered an interpretable breakdown of the contributions of individual metrics, accounting for the shared variance among them.

| **Metric** | ***Percentage Explained (%)*** |
|---|---|
| Average Betweeness Centrality | 2.32 |
| Average Clustering | 77.26 |
| Average Path Lenght | 2.42 |
| Modularity | 18.04 |
| Segregation | 0.01 |
| Degree Standard Deviation | 0.02 |

**Table S2:** Contribution of each variable between Ising Temperature and the graph metrics.

**Acute**

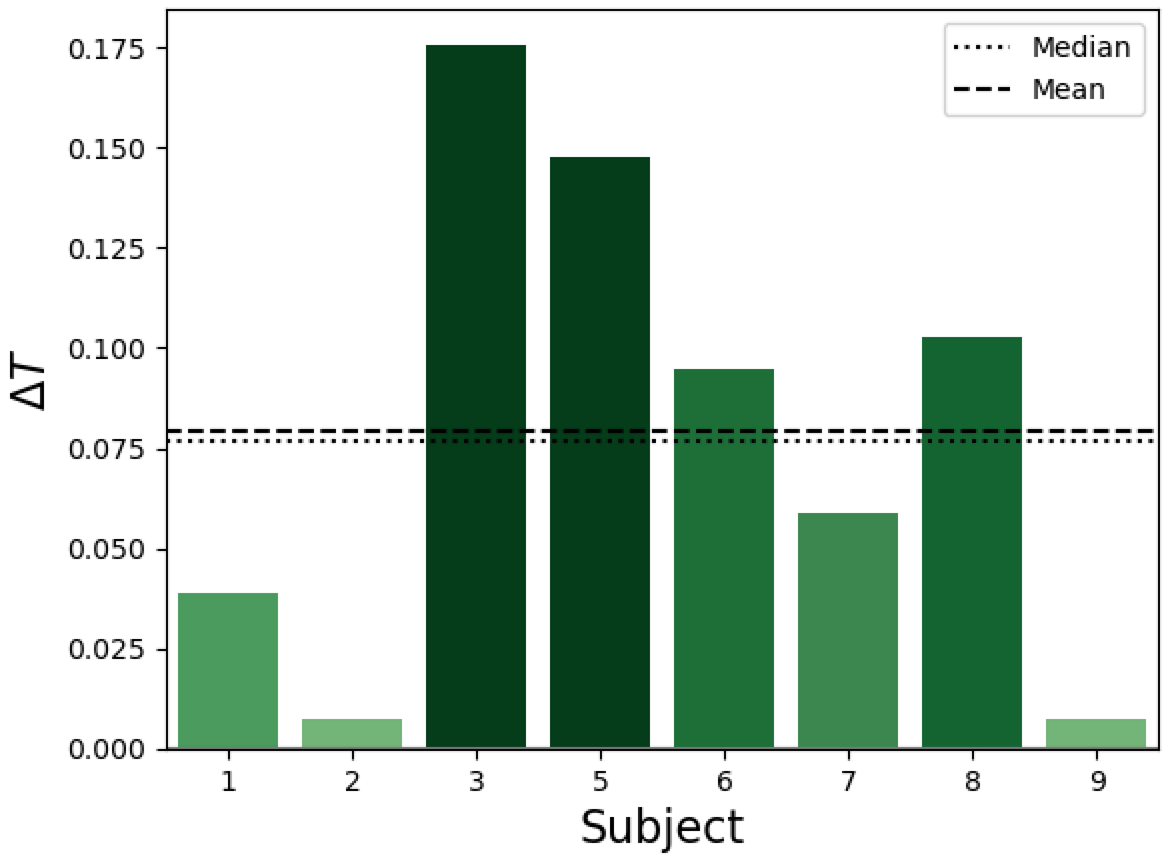


**Figure S1:** Bar chart displaying the individual changes in Ising temperature (ΔT) for each subject in the ayahuasca group during the acute phase. The dashed lines represent the mean and median ΔT values across all subjects. The chart highlights notable variability in temperature changes, with some subjects (e.g., Subjects 3 and 5) showing significant increases compared to others. This variability suggests differential responses in cortical functional connectivity after ayahuasca administration.

**Subacute**

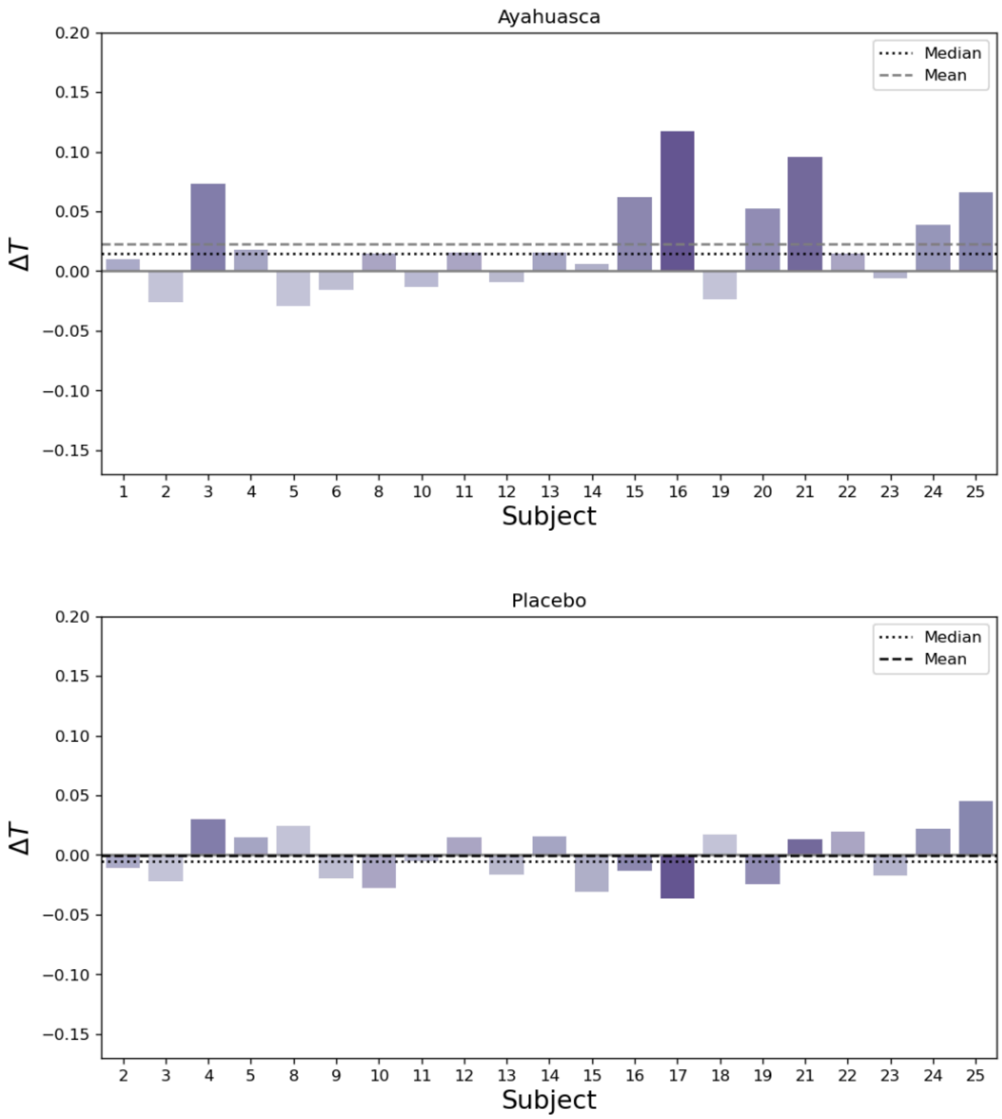


**Figure S2:** Visualization of the changes in Ising temperature (ΔT) for each subject during the subacute phase following ayahuasca and placebo administration. The figure shows a significant increase in Ising temperature in the ayahuasca group compared to the placebo group in absolute values above the mean (dashed line) and median (dotted line).

| | Acute | Subacute |
|---|---|---|
| *Adj. $R^2$* | 0.38 | 0.04 |
| | ***p value*** | |
| Translation X | 0.41 | 0.62 |
| Translation Y | 0.07 | 0.31 |
| Translation Z | 0.55 | 0.88 |
| Rotation X | 0.53 | 0.78 |
| Rotation Y | 0.40 | 0.45 |
| Rotation Z | 0.21 | 0.88 |

**Table S3:** Summary of linear regression (Independent Variables: Average Movement; Dependent Variable: Ising temperature change($\Delta T$)) for potential confounding effects due to head motion during the acute and subacute phases. The table presents R-squared ($R^2$) values and p-values for translational (X, Y, Z) and rotational (X, Y, Z) movements. The low $R^2$ values and non-significant p-values suggest minimal impact of head motion on the observed changes in brain connectivity metrics.

| | Ayahuasca | | | Placebo | | |
|---|---|---|---|---|---|---|
| | Temperature Baseline | Temperature After | Temperature Change | Temperature Baseline | Temperature After | Temperature Change |
| Adj. $R^2$ | -0.19 | -0.14 | 0.58 | -0.09 | -0.52 | -0.05 |
| β Affect average | *p > 0.1* | *p = 0.06* | p < 0.01 | *p > 0.1* | *p > 0.1* | *p > 0.1* |
| β Perception Avg. | *p > 0.1* | *p > 0.1* | *p = 0.06* | *p > 0.1* | *p > 0.1* | *p > 0.1* |
| β Somaesthesia Std | *p > 0.1* | *p > 0.1* | *p = 0.07* | *p > 0.1* | *p > 0.1* | *p > 0.1* |
| β other | *p > 0.1* | *p > 0.1* | *p > 0.1* | *p > 0.1* | *p > 0.1* | *p > 0.1* |

**Table S4:** Summary of linear regression (Dep. Var. HRS and Ind. Var. Ising Temperature) with Adjusted R-squared (Adj. $R^2$) values and p-values for temperature measurements before and after administration, and temperature changes in the ayahuasca and placebo groups. The table shows a significant effect on the affect average ($p < 0.01$) for temperature changes in the ayahuasca group, while no significant changes were observed in the placebo group ($p > 0.1$).
